# Egocentric cues influence the spatial memory of landmark configurations for memory-guided actions

**DOI:** 10.1101/2023.04.11.536430

**Authors:** Pierre-Pascal Forster, Katja Fiehler, Harun Karimpur

## Abstract

Allocentric and egocentric reference frames are used to determine the spatial position of action targets in reference to objects in the environment, i.e., landmarks (allocentric), or the observer (egocentric). Previous research investigated reference frames in isolation, for example, by shifting landmarks relative to the target and asking participants to reach to the remembered target location. Systematic reaching errors were found in the direction of the landmark shift and used as a proxy for allocentric spatial coding. Here we examined the interaction of both allocentric and egocentric reference frames by shifting the landmarks *as well as* the observer. We asked participants to encode a 3D configuration of balls, and to reproduce this configuration from memory after a short delay followed by a landmark or observer shift. We also manipulated the number of landmarks to test its effect on the use of allocentric and egocentric reference frames. Shifting the observer resulted in larger configurational errors. In addition, an increase in the number of landmarks led to a stronger reliance on allocentric cues and a weaker contribution of egocentric cues. In sum, our results highlight the important role of egocentric cues for allocentric spatial coding in the context of memory-guided actions.

**New & Noteworthy:** Objects in our environment are coded relative to each other (allocentrically) and are thought to serve as independent and reliable cues (landmarks) in the context of unreliable egocentric signals. Contrary to this assumption, we demonstrate that egocentric cues alter the spatial memory of landmark configurations, which could reflect recently discovered interactions between allocentric and egocentric neural processing pathways. Further, additional landmarks lead to a higher contribution of allocentric and a lower contribution of egocentric cues.

## Introduction

Different classes of spatial reference frames are used for interacting with objects (1). For example, when we grab a cup of tea, the cup can be represented relative to the observer, i.e., in an egocentric reference frame. Visually-guided actions primarily rely on an egocentric gaze-centered reference frame in which the action target is represented with respect to the current position of gaze (2). As a consequence, continuous spatial updating of the target representation is required to compensate for changes in the spatial origin with an observer’s movement (3).

The cup can also be represented with respect to other objects in the environment, such as the tea pot, i.e., in an allocentric reference frame. Relying on multiple landmarks can improve allocentric spatial coding by providing a more stable point of origin compared to a single landmark (4). As allocentric cues are independent of the observer’s movement, they are more persistent over time (5) and therefore especially suitable for memory-guided movements (6; 7).

The so called “object-shift paradigm” is a prominent task to investigate the contribution of allocentric reference frames for memory-guided actions. In this paradigm, participants are asked to reach to the remembered location of a target object while surrounding objects (landmarks) are unnotably shifted (7; 8). The results show that reach endpoints systematically deviate in the direction of the landmark shift, reflecting the use of allocentric reference frames for memory-guided actions. These deviations are larger if more landmarks are shifted (4; 8). Nevertheless, deviations in reach endpoints are usually only halfway the actual landmark shift, even if all available landmarks are shifted in the same direction (4; 7; 8). This suggests an additional contribution of egocentric reference frames that has never been tested in the “object-shift paradigm”.

Previous research shows that targets for action are concurrently represented in multiple spatial reference frames (9). When allocentric and egocentric cues are available, they seem to be combined by additionally considering the stability of the allocentric cues (10; 11). However, these studies leave open whether and how allocentric and egocentric cues interact and impact each other. The extent to which neural pathways for allocentric and egocentric reference frames are intertwined (9), gives reason to believe that allocentric configurations could be altered by egocentric signals leading to behavioral changes.

In this study, we investigated the interaction of allocentric and egocentric reference frames in an adapted version of the object-shift paradigm. We shifted the landmarks (allocentric cues) *and* the observer (egocentric cues) and varied the number of available landmarks to test its influences on allocentric and egocentric coding. Participants were asked to reproduce the spatial configuration of the landmarks by placing one to five virtual balls after a memory delay and the respective shift at the remembered positions in a virtual environment. We analyzed participants’ placement error relative to the maximum expected error due to the landmark (allocentric weight) or observer shift (egocentric weight).

We hypothesized that placement errors should increase after landmark and observer shifts, if participants rely on allocentric and egocentric cues, respectively. If egocentric cues influence allocentric spatial coding, it should be reflected in decreased allocentric weights or increased configurational errors when the observer is shifted. Further, if the number of landmarks influences the relative contribution of allocentric and egocentric cues, allocentric weights should decrease and egocentric weights increase the fewer landmarks are available. Likewise, configurational errors should increase when fewer landmarks are available.

## Methods

### Participants

Participants were recruited via university e-mail and received either monetary compensation or course credits for their participation. We screened participants for normal stereo vision with the Stereo Fly Test (Stereo Optical Co., Inc., Chicago, USA) and right handedness with the Edinburgh Handedness Inventory (EHI, 12). Three participants failed the criterion for stereo vision (values of ≥6 required), and another three were excluded because their average ball placement errors were greater than the median error across participants plus the median absolute deviation. The final sample consisted of 35 participants (24 female, 11 male) with a mean age of 23.14 (*SD* = 3.50) years. The experiment was approved by the local ethics committee, and is in line with the Declaration of Helsinki (2013, except §35, pre-registration). All participants provided written informed consent.

### Setup and Stimuli

We used a Vive Pro Eye (HTC Corporation, Taoyuan City, Taiwan) head mounted display (HMD, 90Hz frame rate, 1440×1600 pixels per eye, 110° field of view) together with two Vive lighthouses and a Vive controller. The experiment was run in Unity (Unity Technologies, San Francisco, CA, USA). The virtual environment consisted of a pebbly underground and a structured blue-grey wall. Using structured instead of uniform materials is essential to create the optic flow pattern, i.e., to simulate an observer shift through a lateral movement of the wall and the underground (13). The configuration of balls consisted of six white balls with a size of 5 cm in radius and a distance of at least 17 cm to each other. Participants stood approximately 55 cm away from this configuration. The starting position was marked on the virtual floor and a tone was played when participants came too close to the configuration of balls. We used three different configurations, one for each number of landmarks (see Design), to refrain participants from learning the balls’ positions.

### Design

To probe allocentric and egocentric coding we introduced a clearly noticeable lateral shift of the observer by inducing an optic flow pattern (*observer shift*), or, unbeknownst to participants, presented a lateral shift of the landmarks (*landmark shift*), respectively. In the baseline condition, neither a shift of the observer nor the landmark(s) occurred. We also manipulated the number of *landmarks*. This resulted in a 3×3×3 within-subject design with the factors landmark shift (left, none, right), observer shift (left, none, right), and number of landmarks (1, 3, 5). We tested all 27 combinations and repeated each combination in two separate blocks (54 trials in total). The order of trials in each block was randomized.

### Procedure

Participants first performed the stereo vision and handedness tests. Each trial started with the presentation of an encoding scene (5000 ms) which consisted of a configuration of six balls (Figure 1). Then, a mask was shown (200 ms) to prevent after effects, followed by a delay (1800 ms) and a test scene (1000 ms) which consisted of 1, 3, or 5 of the landmarks of the encoding scene. In the test scene, landmarks could be laterally shifted by 4 cm (jittered by ±5 mm). Afterwards, an observer shift of 20 cm could be simulated by a lateral optic flow pattern (1000 ms). After the respective shift(s), participants used the controller to successively place the missing balls (1, 3, or 5) back to their original positions in order to reconstruct the configuration of the encoding scene from memory. No landmarks were present during the observer shift and the ball placement.

**Figure 1.**
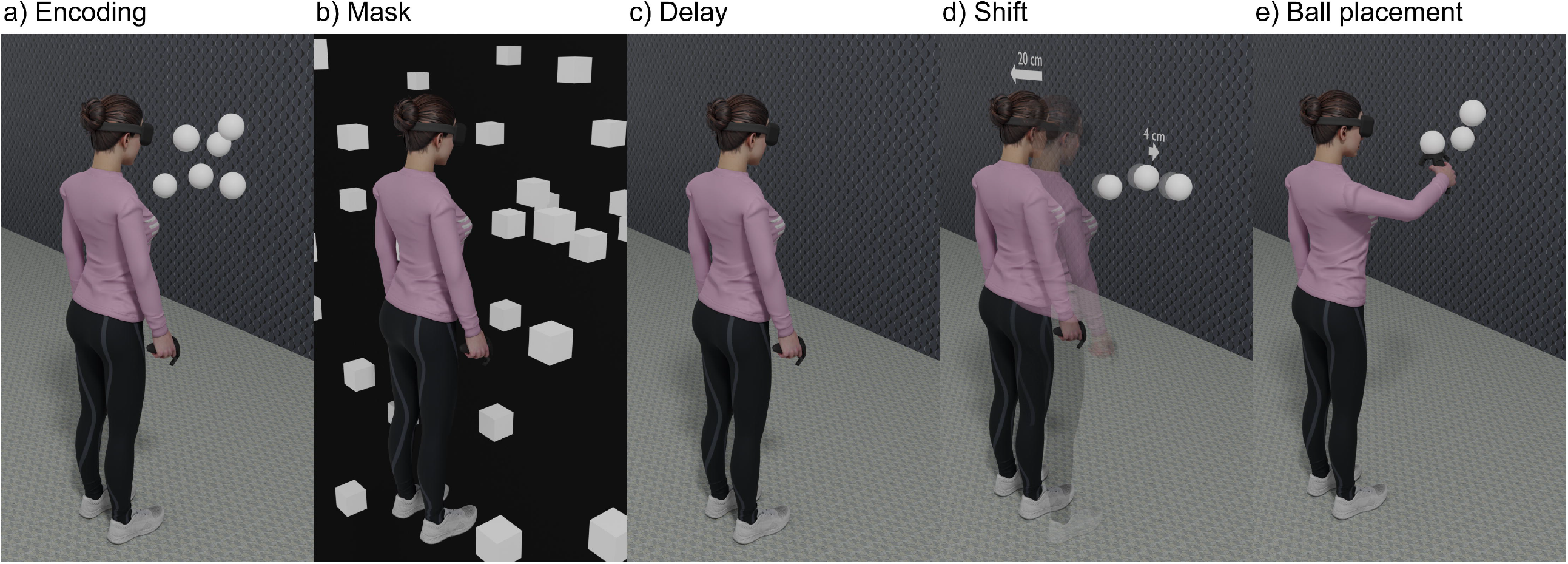
Depiction of a participant performing the experiment. a) A participant encodes the scene in which a configuration of six balls was presented. After b) a mask and c) a brief delay, d) either the landmarks, the participant or both were shifted. If both shifts occurred, the landmarks were presented at their new, shifted location (unnotably to the participant) after the delay, followed by the lateral shift of the participant. e) The participant uses the controller to place the missing balls at the remembered positions in an empty scene. The avatar model was not visible to participants and taken from www.mixamo.com.

### Pre-processing and analysis

We used Python 3.8 to pre-process the data. To compare the contribution of each reference frame independent of the difference in shift amplitude, we calculated *allocentric* and *egocentric weights*. The baseline corrected placement error (PE_corr_) for the allocentric weight was calculated by subtracting the ball positions in the baseline from the respective ball positions for left and right landmark shifts, and subsequently averaging the resultant errors over all ball positions within a given ball configuration (see Figure 2). We then subtracted the (lateral) PE_corr_ for landmark shifts to the right from the ones with shifts towards the left, and divided these relative errors by the difference of the maximum expected placement error (PE_max_) for left and right landmark shifts. The PE_max_ equals the amount of landmark shift, i.e., -4 cm for left and 4 cm for right shifts. The calculation for the egocentric weights was analogous, the only difference being that the weights were calculated with respect to the observer shift (PE_max_ = ±20 cm), instead of the landmark shift. Merging the three respective shift directions reduced the data to 18 data points per participant and weight. Higher weights indicate a stronger reliance on the respective reference frame. For example, an allocentric weight of 0 indicates that participants did not make use of allocentric cues for the ball placement, whereby an allocentric weight of 1 indicates that they solely relied on this cue.

**Figure 2.**
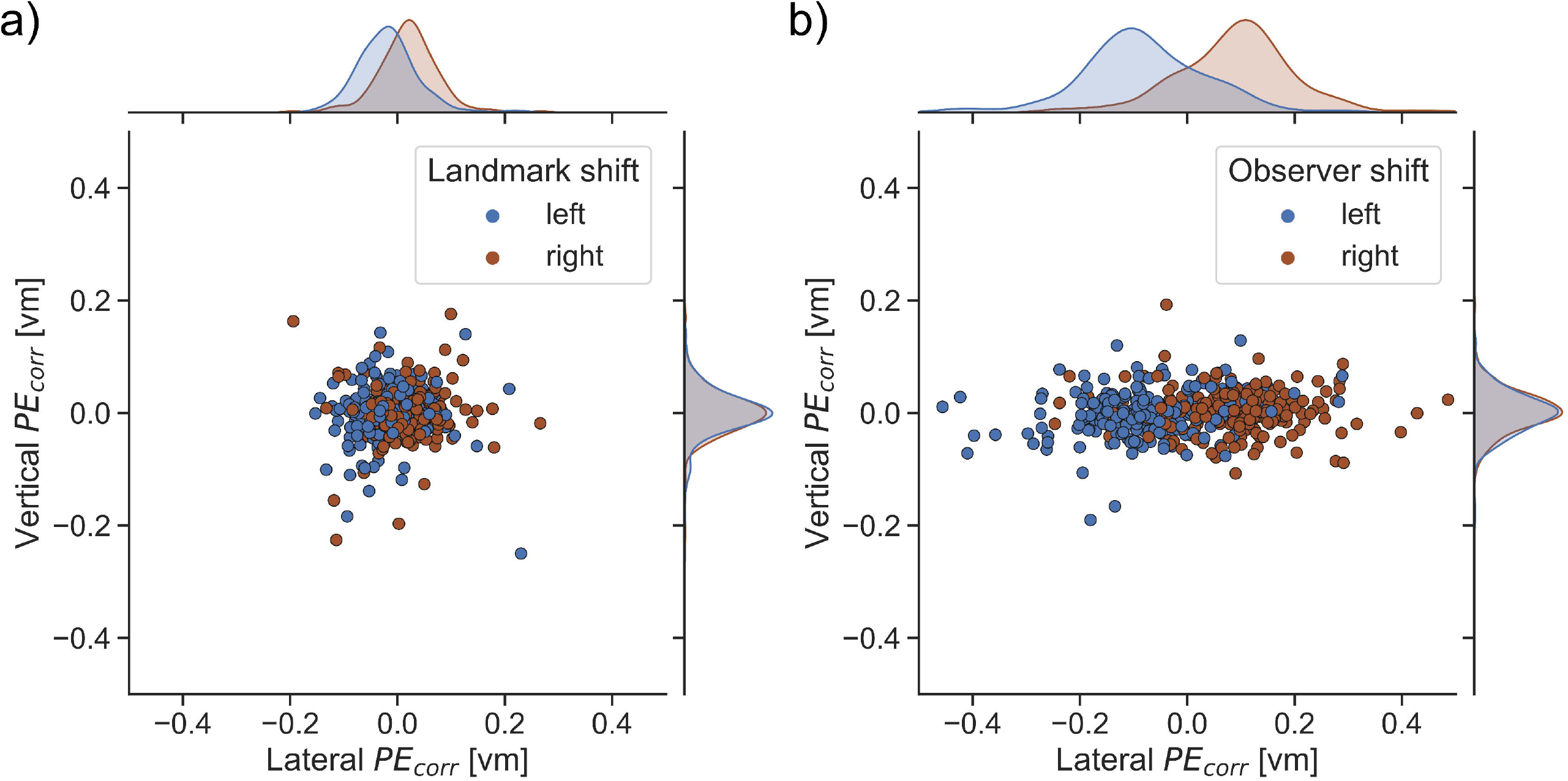
Baseline corrected placement error (PE_corr_) in virtual meters (vm) for a) the effect of a landmark shift when no observer shift was present and b) the effect of an observer shift when no landmark shift was present. Negative values indicate errors towards the left and positive values towards the right.

The *lateral centroid distance* was calculated as the difference of the means between the initial and reproduced configuration for the coordinates along the lateral axis. Negative values indicate a shift of the reproduced configuration towards the left, positive values towards the right. The *configurational error* was calculated as root-mean-square-error (RMSE) of the disparity from the Procrustes analysis (14; 15). It specifies how well the initially presented configuration matches the reproduced configuration of balls after transforming (e.g., rotating and scaling) the input configuration. An error of 0 indicates that the initial and the (transformed) reproduced configuration are identical and higher positive values indicate an increasingly worse spatial overlap. As a configuration requires at least two balls, we calculated the configurational error only for trials where participants placed five and three missing balls, i.e., when one or three landmarks were presented.

We excluded values outside 1.5 times the interquartile range for the allocentric weights (6.03%), egocentric weights (4.29%), lateral centroid distance (3.02%), and configurational error (0.24%). To analyze those data, we used R 4.1.0 to run a linear mixed model (LMM) using the *lmerTest* package (for a primer on LMMs see 16). To account for the within-subject design we added a random intercept per participant to the model (see Supplemental Table S1). We used dummy coding as a coding scheme and Restricted Maximum Likelihood (REML) to fit the models (default in R). For the lateral centroid distance, we used all three levels of landmark and observer shifts in the model to test if those manipulations shifted the centroid in the expected direction. For all other models, observer and landmark shift variables were binarized. Contrasts for post-hoc tests were based on the estimated marginal means from the *emmeans* package with Holm correction applied.

## Results

To investigate the interaction of allocentric and egocentric reference frames, we simultaneously or separately manipulated allocentric and egocentric cues and measured the effect on participants’ ball placement behavior when reproducing the balls’ configuration from memory. Figure 2 shows first descriptive results of the PE_corr_ in lateral and vertical directions. Landmark and observer shifts clearly introduced placement errors in the lateral direction.

### Allocentric weights

The allocentric weights (Figure 3a) differed depending on the number of landmarks in the test scene (*F*(2, 552.893)=5.369, *p*=.005, η²=.018). This effect was influenced by the observer shift (*F*(2, 552.582)=3.483, *p*=.031, η²=.012). Post-hoc tests showed that all allocentric weights differed from zero (all *p*≤.003), indicating the use of allocentric cues when reproducing the spatial configuration of balls. When no observer shift occurred, allocentric weights were lower when one compared to all five landmarks were presented (*t*(552.500)=2.590, *p*=.030). Similarly, when an observer shift occurred, allocentric weights were lower when three compared to five landmarks were presented (*t*(554.896)=3.342, *p*=.003). This indicates that allocentric weights decrease when allocentric cues become less reliable, but this influence varies with the observer shift. For three landmarks, the allocentric weights were smaller when an observer shift occurred compared to no shift (*t*(553.448)=2.097, *p*=.036), demonstrating an influence of egocentric cues on allocentric spatial coding.

**Figure 3.**
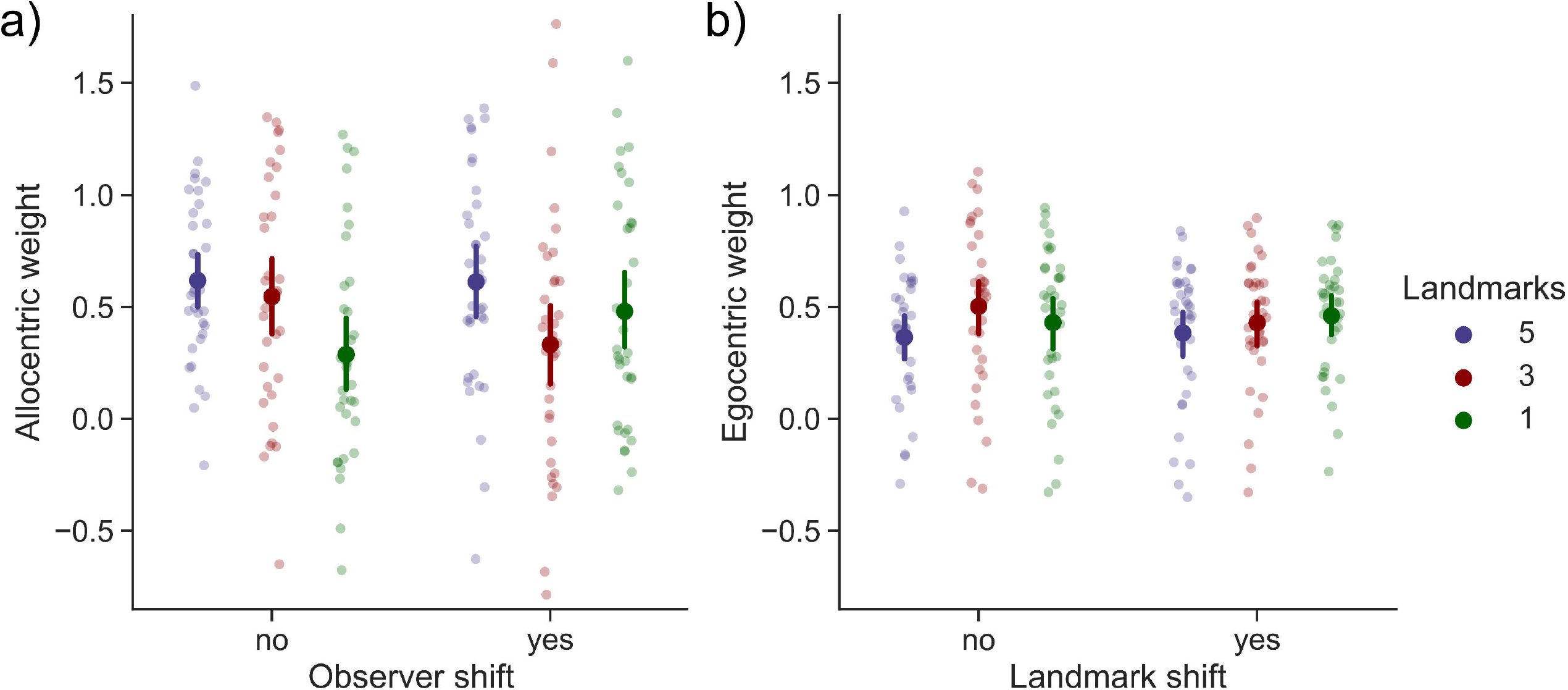
Mean a) allocentric and b) egocentric weights depending on the shift condition and the number of landmarks. Individual data points are averaged per participant. Error bars represent the 95% within-subject confidence intervals. Note that the factor landmark shift was not significant for the egocentric weights, but was kept in the visualization to facilitate comparing allocentric and the egocentric weights.

### Egocentric weights

The egocentric weight (Figure 3b) was influenced by the number of landmarks (*F*(2, 565.237)=5.049, *p*=.007, η²=.017). Compared to five landmarks, conditions with three (*b*=0.073 [CI: 0.024 – 0.121], SE=0.025, *t*(565.233)=2.934, *p*=.003) and one (*b*=0.062 [CI: 0.014 – 0.110], SE=0.025, *t*(565.371)=2.513, *p*=.012) landmarks had a higher egocentric weight. This indicates that the reliance on egocentric cues increases when allocentric cues become less reliable. Post-hoc tests showed that all egocentric weights were different from zero (all *p*<.001), indicating the use of egocentric cues to reproduce the spatial configuration.

### Lateral centroid distance

For the lateral centroid distance (Figure 4a and b), we found an effect of landmark shift (*F*(2, 1788.099)=26.897, *p*<.001, η²=.019). As expected, the centroid was misplaced to the left when the landmarks were shifted to the left (*b*=-0.021 [CI: -0.030 – (−0.012)], SE=0.005, *t*(1788.039)=-4.599, *p*<.001), and to the right when the landmarks were shifted to the right (*b*=0.012 [CI: 0.003 – 0.021], SE=0.005, *t*(1788.139)=2.639, *p*=.008). The influence of landmark shift on the lateral position of the centroid of the balls’ configuration suggests that participants relied on allocentric cues to place the balls at their remembered position.

**Figure 4.**
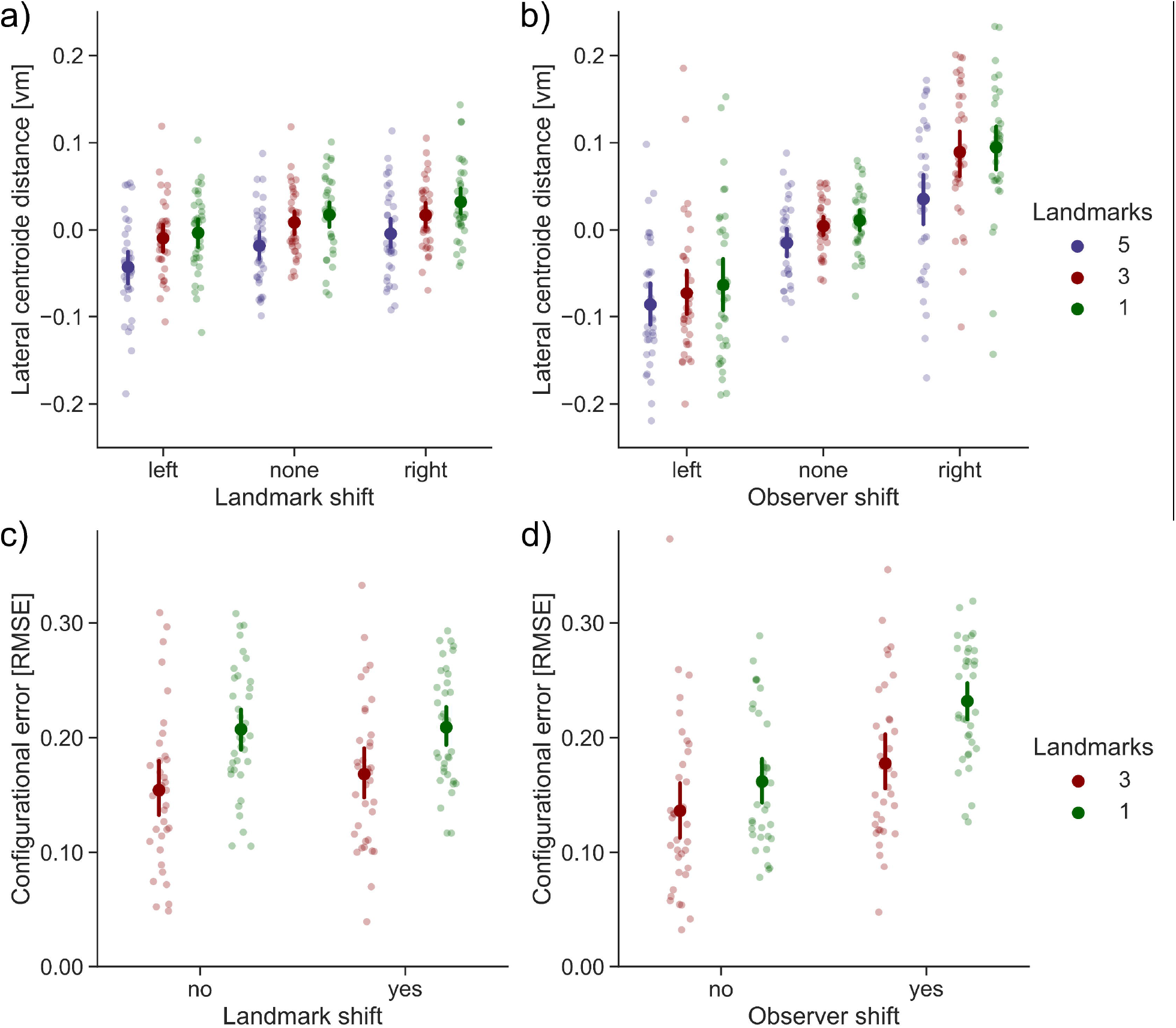
Means for the lateral centroid distance and the configurational error. a) and b) show the lateral centroid distance in virtual meters (vm) for landmark and observer shift, respectively. c) and d) show the configurational error as root-mean-square-error (RMSE) for landmark and observer shift, respectively. Note that the effect of landmark shift for the configurational error is only displayed in the figure for visualization purposes. Individual data points are averaged per participant. Error bars represent the 95% within-subject confidence intervals.

The observer shift (*F*(2, 1789.247)=484.966, *p*<.001, η²=.334) and the number of presented landmarks (*F*(2, 1788.512)=33.437, *p*<.001, η²=.023) interacted with each other (*F*(4, 1788.277)=5.373, *p*<.001, η²=.007). Post-hoc tests showed that, similar to the landmark shift, the centroid positions differed between the different observer shifts (left, none, right) for each number of presented landmarks (all *p*<.001), i.e., participants reproduced the centroid in the direction of the observer shift. This indicates that participants also relied on egocentric cues. Moreover, we found an overall rightward error that depended on the number of landmarks. In detail, the centroid was reproduced farther to the right when only one landmark was presented compared to five landmarks for all three observer shifts (all *p*≤.028). The same was true for three compared to five landmarks when the observer was not shifted (*t*(1788.281)=-2.292, *p*=.044) or shifted towards the right (*t*(1788.644)=-6.585, *p*<.001). Presenting fewer landmarks decreased the effect on the lateral centroid distance if the observer was shifted to the left, and increased the effect if the observer was shifted to the right.

### Configurational error

Shifting the observer compared to no shift increased the configurational error (*b*=0.041 [CI: 0.025 – 0.057], SE=0.008, *t*(1219.009)=4.889, *p*<.001), which indicates that the spatial representation of the landmark configuration is influenced by egocentric cues (Figure 4c and d). In addition, the availability of one landmark in the test scene produced a more pronounced configurational error compared to three landmarks (*b*=0.025 [CI: 0.007 – 0.044], SE=0.010, *t*(1218.970)=2.653, *p*=.009). We also found an interaction between the number of landmarks and the observer shift (*b*=0.029 [CI: 0.006 – 0.052], SE=0.012, *t*(1218.989)=2.446, *p*=.015). The difference in the configurational error between one and three landmarks increased with an additional observer shift. Thus, having to place more balls at the remembered locations (i.e., fewer landmarks) and changing the participants’ position increased the configurational error, especially if both appeared together.

## Discussion

We investigated the interaction of allocentric and egocentric reference frames in an adapted version of the “object-shift paradigm”. We manipulated the spatial location of the landmarks and the observer, and varied the number of allocentric cues. Here we report unprecedented findings of an egocentric influence on allocentric spatial coding for memory-guided actions. In line with previous studies (4; 8), we found decreased reliance on allocentric cues when fewer landmarks were available. More importantly, the spatial configuration (arrangement of balls) was reproduced less accurately when egocentric shifts were induced, suggesting an influence of egocentric cues on the memory of landmark configurations. This challenges the idea that objects that are factually independent of the observer’s own movements are also processed independently.

Two views, one from a theoretical stance and one from a neurophysiological stance, could explain our findings. From a theoretical stance, it has been argued that all spatial reference frames are egocentric (17). The very moment humans encode spatial configurations, they do so from a certain – egocentric – vantage point. It is therefore in the nature of spatial coding itself that egocentric cues have an impact on how humans code spatial configurations. From a neurophysiological stance, findings suggest a more complex interplay between brain regions relevant for allocentric and egocentric coding, respectively (9). While distinct regions are associated with individual reference frames, several areas in the frontal and posterior parietal cortex are associated with both reference frames and serve as conversion areas. The integration of allocentric information into the main egocentric processing pathway is assumed to take place between target representation and action panning – a rather early stage that could explain the diffusion of individual landmark locations and consequently errors when reproducing them (9).

The spatial position of the configuration’s centroid was biased in the direction of the observer or the landmark shift, indicating the use of allocentric and egocentric cues. We also observed a rightward bias of the centroid with fewer available landmarks, possibly due to participants’ right handedness (c.f., 18) and fewer spatial reference points to facilitate memory recall (19). Presenting fewer landmarks likely increases spatial uncertainty which can lead to a stronger reliance on egocentric cues (10).

Accordingly, we observed allocentric and egocentric weights above zero when the landmarks or the observer were shifted, supporting that participants use both reference frames. Importantly, their relative contribution was influenced by the number of available landmarks. When fewer landmarks were presented, participants relied more strongly on egocentric and less on allocentric cues. This is in line with previous studies showing that allocentric weights increase with the number of shifted objects (4; 8).

Shifting the observer reduced the allocentric weights for three landmarks, and descriptively increased the weights for one landmark. This unexpected effect might mainly be driven by higher allocentric weights for one landmark if participants were shifted towards the left (see Supplemental Figure S1). Previous work showed reduced reaching errors when landmarks were viewed in the left peripheral visual field (20). However, in our experiment, simulating an observer shift to the left resulted in moving the *remembered* target positions towards participants’ right, where no such effect was observed. Other spatial biases, such as biased attention towards the left (21; 22), biased peripersonal space perception (23), or participant’s handedness (c.f., 18), were controlled by using baseline corrected errors when calculating allocentric weights. Perceiving movements towards the left as faster (24), might cause participants to place the missing balls further towards the right. However, this effect should be cancelled out by combining placement errors for left-and rightward landmark shifts.

In summary, our results show that egocentric cues influence the spatial memory of landmark configurations. The relative contribution of allocentric and egocentric reference frames seems to depend on the number of allocentric cues, with stronger allocentric coding when more landmarks are available. Thus, allocentric and egocentric reference frames do closely interact in memory-guided actions.

## Acknowledgements

We would like to thank Yanina Tena Garcia for her support in the data collection process. The work presented here was funded by the German Research Foundation, International Research Training Group, IRTG 1901, “The Brain in Action” and the DFG grant FI 1567/6-1 “The active observer”.

