## Supplemental Table S1 and Figure S1. for "Egocentric cues influence the spatial memory of landmark configurations for memory-guided actions"


| Model | *b* | CI | SE | *t* | *df* | *p* |
|---|---|---|---|---|---|---|
| Allocentric weights | 0.622 | 0.432 – 0.812 | 0.097 | 6.386 | 396.667 | <.001 |
| Egocentric weights | 0.379 | 0.284 – 0.475 | 0.048 | 7.862 | 39.429 | <.001 |
| Centroid Distance | -0.011 | -0.026 – 0.004 | 0.008 | -1.376 | 198.690 | =.170 |
| Configurational Error | 0.136 | 0.117 – 0.155 | 0.010 | 14.051 | 97.384 | <.001 |

Table S1. Intercepts for the different linear mixed models. Columns depict the slope (*b*), 95% confidence interval (CI), standard error (SE), *t*-value (*t*), degrees of freedom (df) and the *p*-value (*p*).

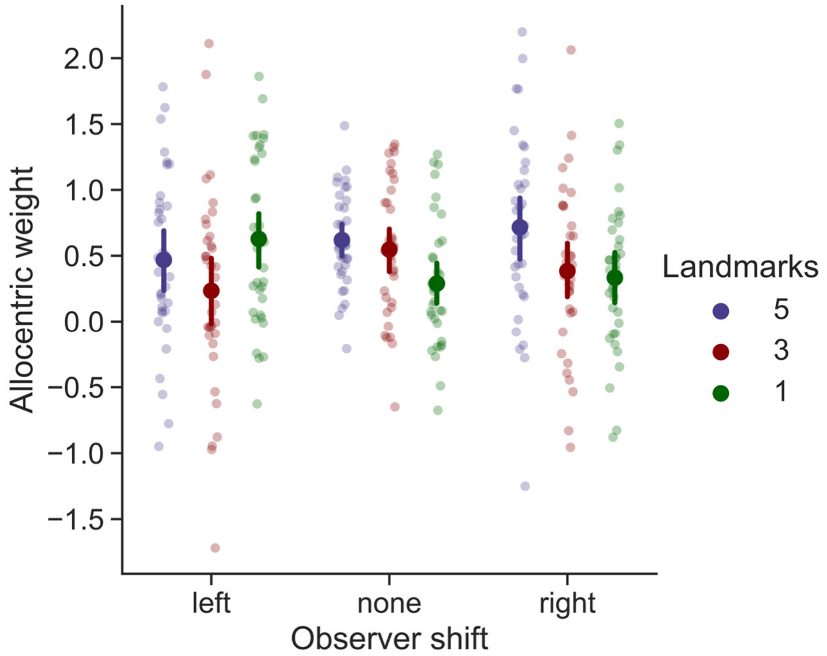


Figure S1. Allocentric weight split over all three observer shifts. Data points are averaged over trial repetitions. All error bars represent the 95% within-subject confidence intervals.
